# A Deep-learning based RNA-seq Germline Variant Caller

**DOI:** 10.1101/2022.10.16.512451

**Authors:** Daniel E. Cook, Aarti Venkat, Dennis Yelizarov, Yannick Pouliot, Pi-Chuan Chang, Andrew Carroll, Francisco M. De La Vega

## Abstract

RNA sequencing (RNA-seq) can be applied to diverse tasks including quantifying gene expression, discovering quantitative trait loci, and identifying gene fusion events. Although RNA-seq can detect germline variants, the complexities of variable transcript abundance, target capture, and amplification introduce challenging sources of error. Here, we extend DeepVariant, a deep-learning based variant caller, to learn and account for the unique challenges presented by RNA-seq data. Our DeepVariant RNA-seq model produces highly accurate variant calls from RNA-sequencing data, and outperforms existing approaches such as Platypus and GATK. We examine factors that influence accuracy, how our model addresses RNA editing events, and how additional thresholding can be used to facilitate our models’ use in a production pipeline.

## Background

RNA-seq is a widely used method for transcriptome analysis. The technology is often used to study gene expression in a variety of contexts. Expression can be examined over time^1^, in response to disease or environmental changes^2,3^, or to examine differences across cell types, tissues, or species^4–7^. In addition to gene expression, RNA-seq can be used for identifying quantitative trait loci (QTL)^8^, quantifying isoform and allele-specific expression^9–11^, detecting gene fusion events^12^, and in identification of RNA-editing events^13,14^. Interestingly, it is also possible to use RNA-seq to perform germline variant calling^15^. However, although germline variant calling with RNA-seq data could be beneficial in many studies, it is an underutilized application of RNA-sequencing^16^.

The lack of studies using RNA-seq for germline variant calling is likely driven by a number of challenges not present with traditional DNA sequencing (DNA-seq) approaches. For example, alignment coverage, a proxy for gene expression, exhibits more dynamic range in RNA-seq than DNA-seq. Coverage can range from little to none to excessively high levels. Low coverage can reduce or eliminate evidence of germline variants and complicate genotype classification^17,18^. Additionally, RNA splicing, allele-specific expression, and RNA-editing can skew allele frequencies, introduce artifacts, or remove variant signals. Sample preparation can also introduce errors. For example, PCR can result in skewed allele frequencies due to amplification bias^19^. Finally, contamination by RNAse enzymes can degrade RNA, and RNA-seq often requires an additional reverse transcription step to produce a cDNA library that can introduce errors^20^. These issues can be challenging to address in clinical and research settings.

Despite these challenges, calling germline variants from RNA-seq data provides a number of advantages over traditional DNA-seq approaches. Foremost is an economic consideration: RNA-seq can provide germline variant calls where one might otherwise sequence an additional DNA sample^21^. This is especially useful when characterizing coding variation which is captured with most RNA-seq approaches, providing a rapid and cost-effective way to interrogate coding regions. Additionally, the versatility of RNA-seq allows germline variation data to be combined with information about transcription and enables QTL discovery^22^.

Existing RNA-seq variant callers use statistical or algorithmic approaches^23,24^. However, these methods are challenging to implement given the numerous additional sources of error and uncertainty in RNA-seq data, and the complex interactions between them. By contrast, deep learning approaches are capable of learning distinct patterns, extracted from sequence data, that are associated with the validity of candidate variants. In recent years, deep learning approaches have been shown to be highly competitive and often exceed the performance of statistical and algorithmic approaches^25^. Here we introduce a new method for performing variant calling on RNA-seq data. We adapt DeepVariant, a deep-learning based variant caller, for use in calling germline variants from RNA-seq data. Our best performing DeepVariant RNA-seq model is trained using data from the Genotype-Tissue Expression (GTEx) consortium,^4,26^ and can overcome challenges associated with RNA-seq variant calling and accurately classify germline variants. Our GTEx-based DeepVariant RNA-seq model outperforms existing approaches, achieving an F1-Score of 0.933 on coding sequences (CDS) and 0.921 on exonic sequences.

The DeepVariant RNA-seq model and code are available under an open-source license at https://github.com/google/deepvariant.

## Methods

### Software Implementation

DeepVariant is a deep-learning based variant caller that uses a convolutional neural network to classify the genotype of a position in the genome by learning how read data correlates with real and spurious variants^27^. To classify variants, DeepVariant first scans a BAM file to identify evidence of SNP or INDEL variation. For example, a candidate variant SNP is identified when two or more bases differ from the reference at a particular locus. When candidate sites are identified, DeepVariant will proceed to generate an example, or model input. Examples are constructed using read pileups by extracting a series of sequence features into channels. Each channel represents a sequence feature such as base, base quality, or mapping quality. These channels are stacked to form a model input for training or inference. Following training, DeepVariant can use the resulting model to classify candidate variants as being homozygous-REF, heterozygous, or homozygous-ALT.

We adapted the data preprocessing stage of DeepVariant to allow for processing of RNA-seq alignment data (BAMs). RNA-seq BAMs, in contrast to DNA-seq BAMs, contain SKIP CIGAR operations which are used to represent skipped regions (**Supplementary Figure S1**) in the reference genome. These are generally caused by splicing events, although structural variation and mismapping of read segments are other possible explanations. Previously, DeepVariant would incorporate these SKIP operations during the local realignment step, resulting in a substantial increase in the computation time. To address this issue, we introduced a new flag (`--split_skip_reads`) that splits reads into multiple segments by removing their SKIP operations. This change allowed DeepVariant to process RNA-seq data efficiently (**Supplementary Figure S2**). With these changes in place, model training and inference operate similar to existing approaches that use DNA-seq.

### Model Training and Evaluation

Once we updated DeepVariant to process RNA-seq data, we trained models using two datasets. For the first model, we used samples from the Genome in a Bottle consortium (GIAB)^27–29^. The GIAB consortium has developed a high quality set of genotypes for seven human cell-lines, derived from two sets of trios and a singleton. These genotypes can be used as training labels or for benchmarking purposes. To train the DeepVariant RNA-seq GIAB model (“DV RNA-seq [GIAB]”), we extracted RNA from cell pellets of lymphoblastoid lines HG001 (GM12878; 10 replicates), and HG002 (GM26105; 3 replicates) obtained from the Coriell institute. This RNA was sequenced with the Tempus xT assay, which includes exome-capture regions that cover the exons of 19,433 human genes (34 Mb target region of the human genome)^30^. Sequenced reads were aligned using STAR v2.5.4. The resulting BAMs were used for training, using data from chr2-19; chr21 and chr22 were used for hyper-parameter optimization, with chr1 and chr20 held out for testing. For training we used the variants from the truth set from Genome-in-a-Bottle v4.1 and restricting to high confidence regions^31^.

We used RNA-seq data from GTEx V8 (dbGaP phs000424.v8.p2) to train our second model (“DV RNA-seq [GTEx]”). The GTEx consortium has extensively characterized human tissue samples from postmortem donors, and provides whole-genome DNA-seq and RNA-seq from a large collection of tissues. RNA-seq data from the GTEx project was sequenced using the Illumina TruSeq non-stranded polyA+ selection protocol. We assigned GTEx donors to a train or test group, and randomly selected 1000 RNA-seq samples from our train donor group (526 unique donors across 41 tissue types; **Supplementary Table S1**). Similarly, we randomly selected 200 RNA-seq samples for testing (74 unique donors across 36 tissue types; **Supplementary Table S1**).

To derive a truth dataset for training the GTEx model, we ran DeepVariant’s highly-accurate WGS model on GTEx DNA-seq samples and used the resulting genotypes as labels for training and testing our RNA-seq model. We refer to this training data as a ‘silver truth’ dataset because our labels are unlikely to be as accurate as GIAB for training. Nevertheless, this approach generated over 80.4 million training examples, 314x more than was used for the GIAB dataset, and provided a set of training examples with more diversity across tissue and ancestry. We also partitioned the genomes in our silver truth dataset: chr1-19 were used for training, chr21 and chr22 were used for tuning, and chr20 was held out for testing. We performed training by subsetting on exonic regions, and by warmstarting from an existing whole-exome sequence trained DeepVariant model. A summary table of the datasets for both the GTEx and GIAB models is provided in **Supplementary Table S2**. We used pre-aligned BAMs and CRAMs provided by the GTEx consortium. The V8 release RNA-seq BAMs aligned using STAR v2.5.3a, and DNA-seq BAMs aligned using BWA (v0.7.13-r1126).

Separately, we also sequenced HG002 (GM24385, GM26105, GM27730) and HG005 (GM26107) on the NovaSeq platform using an mRNA poly-A based library preparation. These samples were processed using the nf-core/rnaseq pipeline^32^ (10.5281/zenodo.1400710). We used HG002 with our GTEx model to establish filtering cutoff thresholds. HG005, which was held out from training for both GIAB and GTEx models, was used to compare performance of both models. Benchmarking was performed using GIAB Truth set v4.2.1. We used hap.py with the RTG vcfeval engine to perform benchmarking (https://github.com/Illumina/hap.py). Variants were required to have a minimum depth of 3x in the RNA-seq sample in order to be considered.

## Results

### DeepVariant RNA-seq Produces Accurate Germline Variant Calls

We trained two DeepVariant RNA-seq models: “DV RNA-seq [GTEx]” and “DV RNA-seq [GIAB]”. We compared the performance of the GTEx and GIAB models using HG005 (GM26107 poly-A capture), which was held out from training for both models. Performance was stratified across coding sequences (CDS), exon, transcript, gene, high-confidence GIAB regions (GIAB-hc), chr20 (which was held out from all training data), and regions with a minimum of 3x depth. The GTEx model outperforms the GIAB model, except for INDEL exon regions (**Table 1**). We chose to focus most of our subsequent analysis on DV RNA-seq [GTEx] given that it outperformed the GIAB model and fits a desirable use case of calling SNPs in CDS regions.

**Table 1:** Performance of GTEx and GIAB trained models on HG005. F1 scores are shown for DV RNA-seq [GTEx] and DV RNA-seq [GIAB] for sample HG005 across different regions. All regions are intersected with a 3x minimum coverage bed file from the HG005 RNA-seq sample. Bolded numbers indicate the best performing model by region based on F1-score for each variant type (SNP or INDEL).

| Model | CDS | Exon | Transcript | Gene | GIAB-hc | chr20 | 3x |
| --- | --- | --- | --- | --- | --- | --- | --- |
| **SNP** |  |  |  |  |  |  |  |
| DV RNA-seq [GTEx] | **0.934** | **0.918** | **0.835** | **0.831** | **0.857** | **0.837** | **0.82** |
| DV RNA-seq [GIAB] | 0.925 | 0.832 | 0.719 | 0.713 | 0.734 | 0.719 | 0.702 |
| **INDEL** |  |  |  |  |  |  |  |
| DV RNA-seq [GTEx] | **0.692** | 0.646 | **0.557** | **0.557** | **0.663** | **0.564** | **0.537** |
| DV RNA-seq [GIAB] | 0.676 | **0.66** | 0.541 | 0.541 | 0.624 | 0.53 | 0.527 |

### Comparison of DeepVariant RNA-seq to other open source callers

We compared the performance of our GTEx and GIAB models with several other RNA-seq variant calling approaches using our GTEx-derived “silver truth” dataset. Our comparison includes the warm-start model trained on whole-exome data and used for training (DV WES; DeepVariant v1.1.0) to see how much training on RNA-seq data improved our model. We also tested GATK HaplotypeCaller (v4.2.4.0)^24^ based on a RNA-seq [workflow developed](https://gatk.broadinstitute.org/hc/en-us/articles/360035531192-RNAseq-short-variant-discovery-SNPs-Indels-) by the GATK team. Finally, we tested Platypus (v0.8.1.2)^33^. Platypus requires the BAM input to be preprocessed with a tool called Opossum (v1.0), which performs filtering and processing of RNA-seq BAMs to optimize them for RNA-seq variant calling^23^. We also tested other callers with and without Opossum-pre-processed BAMs.

We used our GTEx test data to compare performance across variant callers. Similar to our model comparison analysis, we stratified across several regions intersected at a 3x minimum coverage. We observe that DV RNA-seq [GTEx] achieves the highest median F1 score across all region stratifications (**Table 2**), performing the best overall in CDS regions (**Figure 1**). We observe the same trend with INDELs as well (**Supplementary Figure S3**). Interestingly, we found that preprocessing with Opossum lowered the performance of DeepVariant, likely because the alignment data was substantially modified from our training dataset. However, our modifications to DeepVariant obviate the need for preprocessing while still resulting in more accurate results.

**Figure 1:**
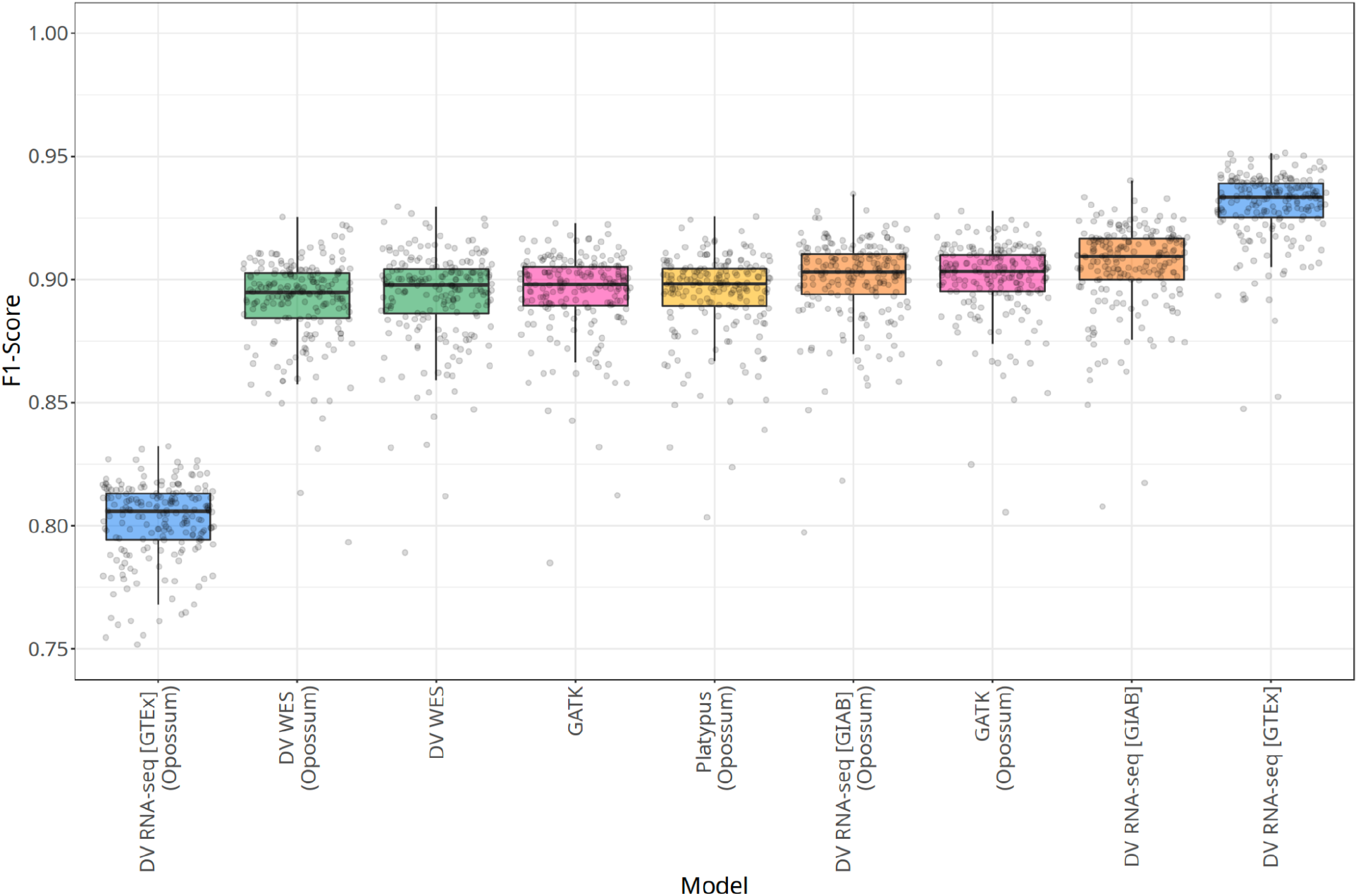
SNP F1 Scores in CDS regions for 200 GTEx RNA-seq samples. SNP F1 scores in CDS regions are shown for 200 GTEx RNA-seq test samples across Platypus, GATK, DV WES, DV RNA-seq [GIAB], and DV RNA-seq [GTEx]. Data that was pre-processed with Opossum is labeled as such.

**Table 2:** Performance of variant callers on 200 GTEx samples across genomic regions. Each row lists the median F1 score across 200 GTEx test samples for the given variant caller when using the original BAM or an Opossum pre-processed BAM in each region type. Performance is sorted by F1-score descending. Bolded numbers indicate the best performing model by region based on F1-score for each variant type (SNP or INDEL).

| Variant Caller | CDS | Exon | Transcript | Gene | chr20 | 3x |
| --- | --- | --- | --- | --- | --- | --- |
| **SNP** |  |  |  |  |  |  |
| DV RNA-seq [GTEx] | **0.933** | **0.921** | **0.846** | **0.842** | **0.83** | **0.83** |
| DV RNA-seq [GIAB] | 0.909 | 0.845 | 0.745 | 0.737 | 0.738 | 0.722 |
| DV RNA-seq [GIAB] (Opossum) | 0.903 | 0.84 | 0.742 | 0.733 | 0.739 | 0.719 |
| GATK (Opossum) | 0.903 | 0.852 | 0.759 | 0.75 | 0.753 | 0.737 |
| DV WES | 0.898 | 0.801 | 0.678 | 0.667 | 0.661 | 0.649 |
| GATK | 0.898 | 0.847 | 0.751 | 0.742 | 0.741 | 0.727 |
| Platypus (Opossum) | 0.898 | 0.833 | 0.726 | 0.717 | 0.716 | 0.703 |
| DV WES (Opossum) | 0.895 | 0.809 | 0.692 | 0.682 | 0.679 | 0.666 |
| DV RNA-seq [GTEx] (Opossum) | 0.805 | 0.785 | 0.701 | 0.696 | 0.687 | 0.685 |
| **INDEL** |  |  |  |  |  |  |
| DV RNA-seq [GTEx] | **0.73** | **0.675** | **0.558** | **0.554** | **0.544** | **0.539** |
| DV RNA-seq [GIAB] | 0.652 | 0.6 | 0.494 | 0.49 | 0.49 | 0.477 |
| DV RNA-seq [GIAB] (Opossum) | 0.592 | 0.558 | 0.436 | 0.432 | 0.443 | 0.418 |
| DV RNA-seq [GTEx] (Opossum) | 0.524 | 0.531 | 0.437 | 0.434 | 0.429 | 0.419 |
| GATK (Opossum) | 0.504 | 0.565 | 0.482 | 0.477 | 0.487 | 0.463 |
| GATK | 0.486 | 0.574 | 0.491 | 0.487 | 0.495 | 0.471 |
| DV WES | 0.423 | 0.581 | 0.493 | 0.489 | 0.502 | 0.477 |
| Platypus (Opossum) | 0.414 | 0.468 | 0.398 | 0.394 | 0.397 | 0.374 |
| DV WES (Opossum) | 0.392 | 0.533 | 0.44 | 0.435 | 0.452 | 0.423 |

We also evaluated the performance on chr20 which was held out from training and again found DV RNA-seq [GTEx] to outperform alternative variant callers (**Supplementary Table S2**).

### Factors Impacting RNA-seq Model Performance

Next, we investigated factors that impact DV RNA-seq [GTEx] performance, focusing on the use case of calling SNPs in CDS regions. We examined F1 scores after restricting variant calling to sites at a series of coverage thresholds in CDS regions (**Figure 2A**). For each threshold we indicate the relative precision and sensitivity.

**Figure 2:**
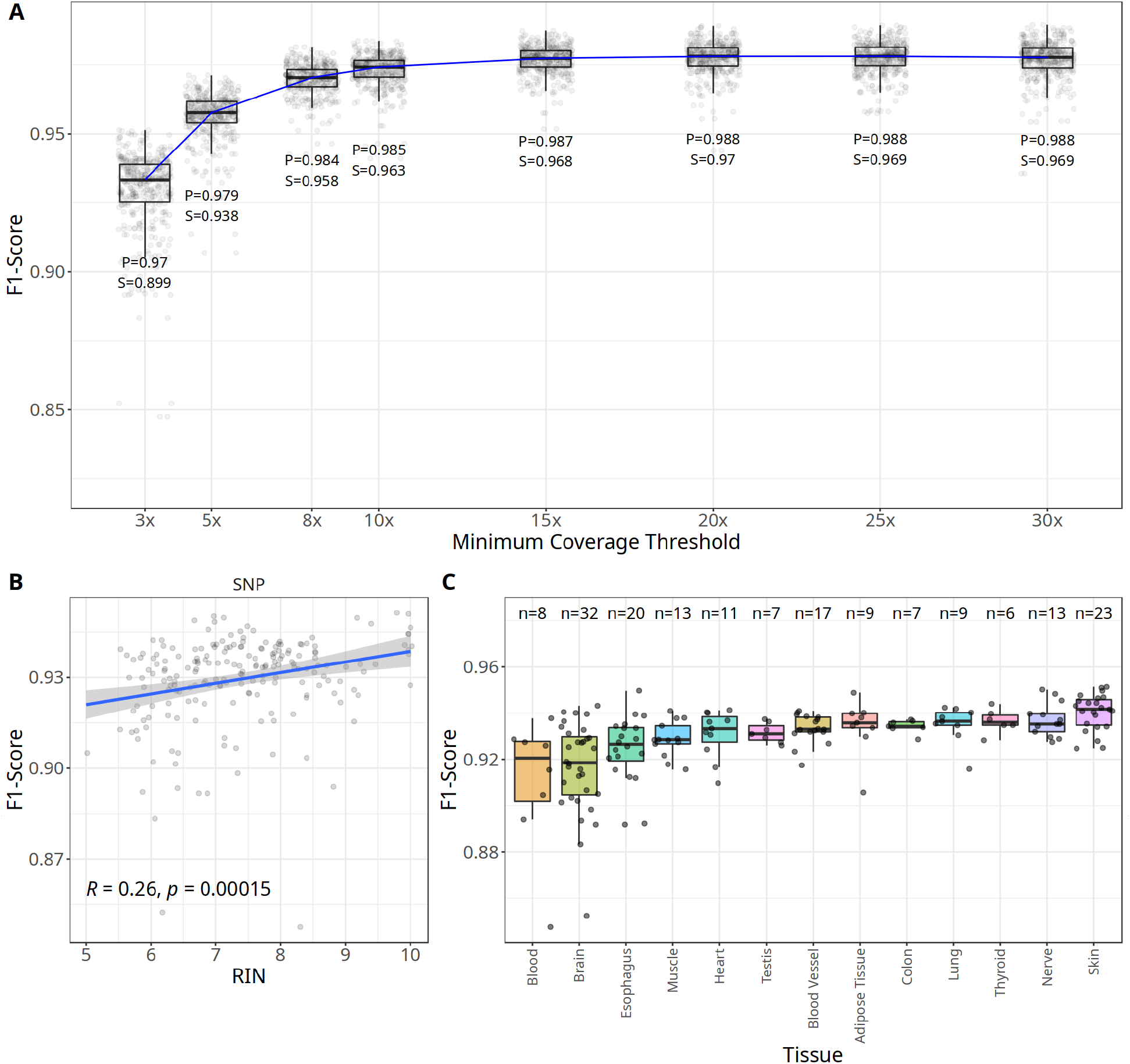
Impact of depth, RIN, and tissue on DV RNA-seq performance. **(A)** F1-scores are shown for SNPs in CDS regions for a series of coverage thresholds. For each coverage threshold, we show the distribution of F1 scores for 200 test RNA-seq samples. Below each box plot are the values for precision (P), sensitivity (S) at the given threshold. **(B)** Correlation plot between RIN and F1-scores for CDS SNPs. **(C)** F1-score distributions for SNPs in CDS regions across a collection of GTEx tissues. The number of RNA-seq samples is indicated for each tissue at the top (n). We filtered tissues with less than 5 samples in our test set.

Assuming variants are detectable at a minimum depth of 3x, we asked what proportion of these detectable variants remain across minimum coverage thresholds (**SupplementaryFigure S4**). Although our examination here will be dependent on coverage for a given sample, our analysis suggests that a modest coverage threshold of 8x sample still captures 75% of CDS label variants on average in GTEx samples.

We also examined the correlation between the RNA Integrity Number (RIN) and RNA-seq sample accuracy. A RIN score can be calculated for a given RNA-sample using an algorithm that examines the ratio of 28S and 18S ribosomal RNA. RIN scores range from 1 (totally degraded) to 10 (intact)^20^, and are used as an indicator of RNA quality. We observe a weak correlation between RIN and CDS SNP F1 scores (R = 0.26; p=0.00015;**Figure 2B**), suggesting that DeepVariant RNA-seq can be used on samples with low RIN scores. We also observe a weak correlation for INDELs as well (R=0.16, p=0.021; **Supplementary Figure S5**).

Tissue type can strongly influence RNA-seq data. We examined how GTEx tissues impacted DeepVariant RNA-seq performance in CDS regions. We observed the highest F1 score with skin and the lowest performance in blood (**Figure 2C**). However, performance for SNPs in CDS regions was relatively consistent across tissues. One major factor that may drive differences in performance across tissue is the proportion of abundant versus rare transcripts, which in turn affects the coverage at variant positions.

Unlike DNA, RNA-seq reads possess splicing boundaries. We reasoned that error rates might increase near exon boundaries where read context around candidate variants will drop off and make accurate variant classification more difficult. Unsurprisingly, we observe this effect across several RNA-seq variant calling methods (**Figure 3**). However, DV RNA-seq [GTEx] appears to largely eliminate this effect for false-positive (FP) SNP variants at exon boundaries, and achieves low false-negative (FN) levels as well.

**Figure 3:**
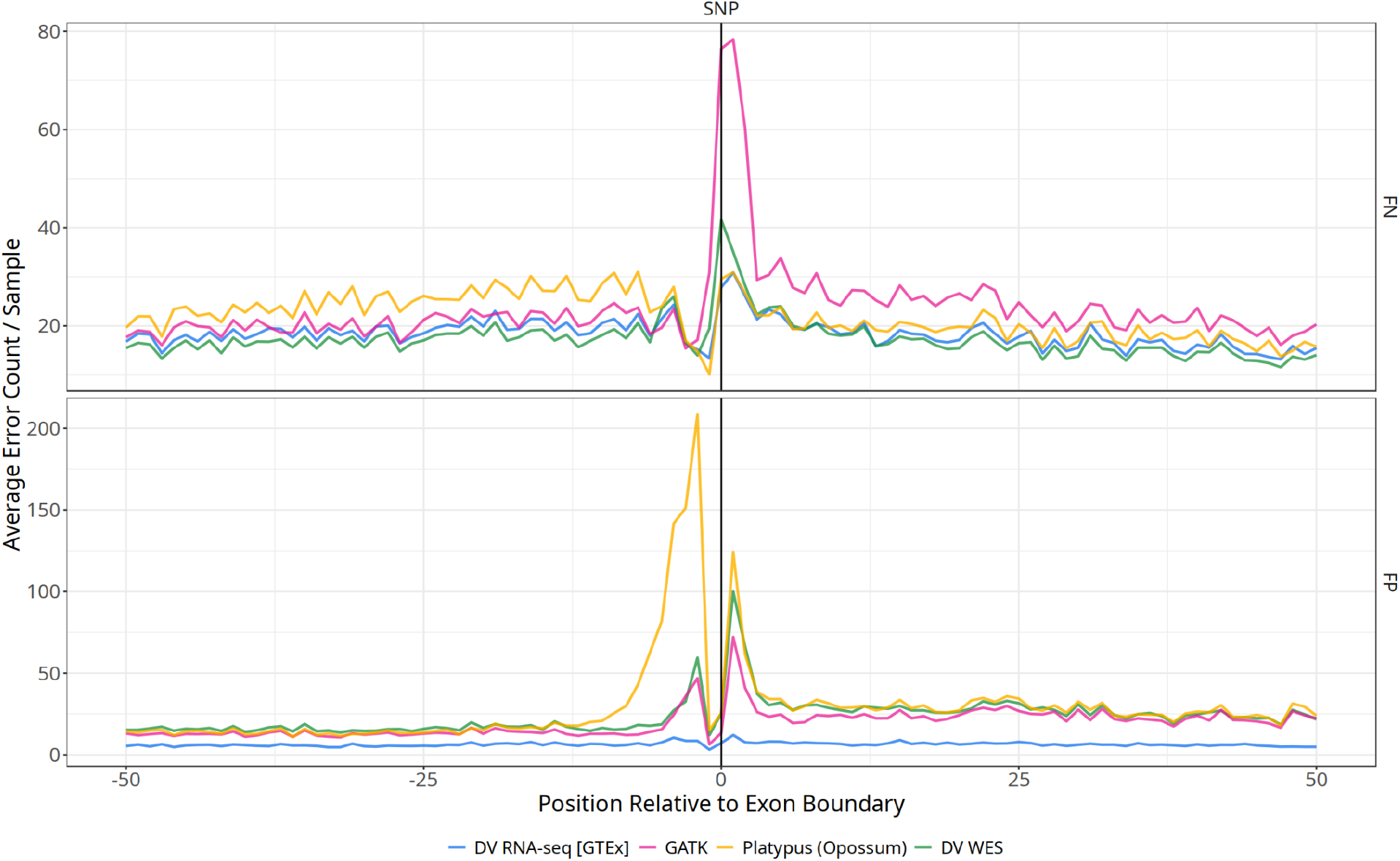
SNP errors relative to exon boundary. Average counts of false negative (FN; top panel) and false positive (FP; bottom panel) errors per GTEx RNA sample (n=200) are shown based on a called SNPs position relative to the nearest exon boundary for several variant callers. Negative positions correspond to SNPs outside of exons, whereas positive positions indicate SNPs falling within exons. A value of 0 indicates a position falling on an exon boundary.

### Our Model Learns to Ignore RNA-editing events

RNA is subject to editing by ADAR (adenosine deaminases acting on RNA), which can result in Adenosine-to-inosine (A-to-I) changes^34^. These A-to-I changes are observed in RNA-seq datasets as apparent A-to-G and T-to-C changes which do not reflect real variants in the DNA, and can increase the germline false positive rate. We sought to examine how our GTEx-based model treated these types of events. To do this, we annotated variants that belonged to an RNA editing database called REDIportal^35^. REDIportal has identified and annotated genomic positions where RNA-editing has been observed using GTEx data.

We asked how the variants compared with the label dataset based on the base change observed and whether they were present in the REDIportal database. Interestingly, we observe that the DV RNA-seq [GTEx] model has a large reduction in the number of false positives at REDIportal sites compared to the DV WES model. In **Figure 4**, we show a representative example of this for GTEx sample GTEX-ZZ64-1126-SM-5GZXY. We also observe this phenomenon in an aggregate analysis (**Supplementary Figure S6**). The REDIportal ~ FP panel shows a considerable reduction in the false-positive rate at sites where RNA editing has been observed. The reduction in false positives does appear to slightly increase the false negative rate at these sites. However, we observe an overall reduction in errors, suggesting that our model learns to ignore RNA-edited sites. We suspect that our model picks up on sequence context and allele frequency differences at these sites as a basis for filtering them out.

**Figure 4:**
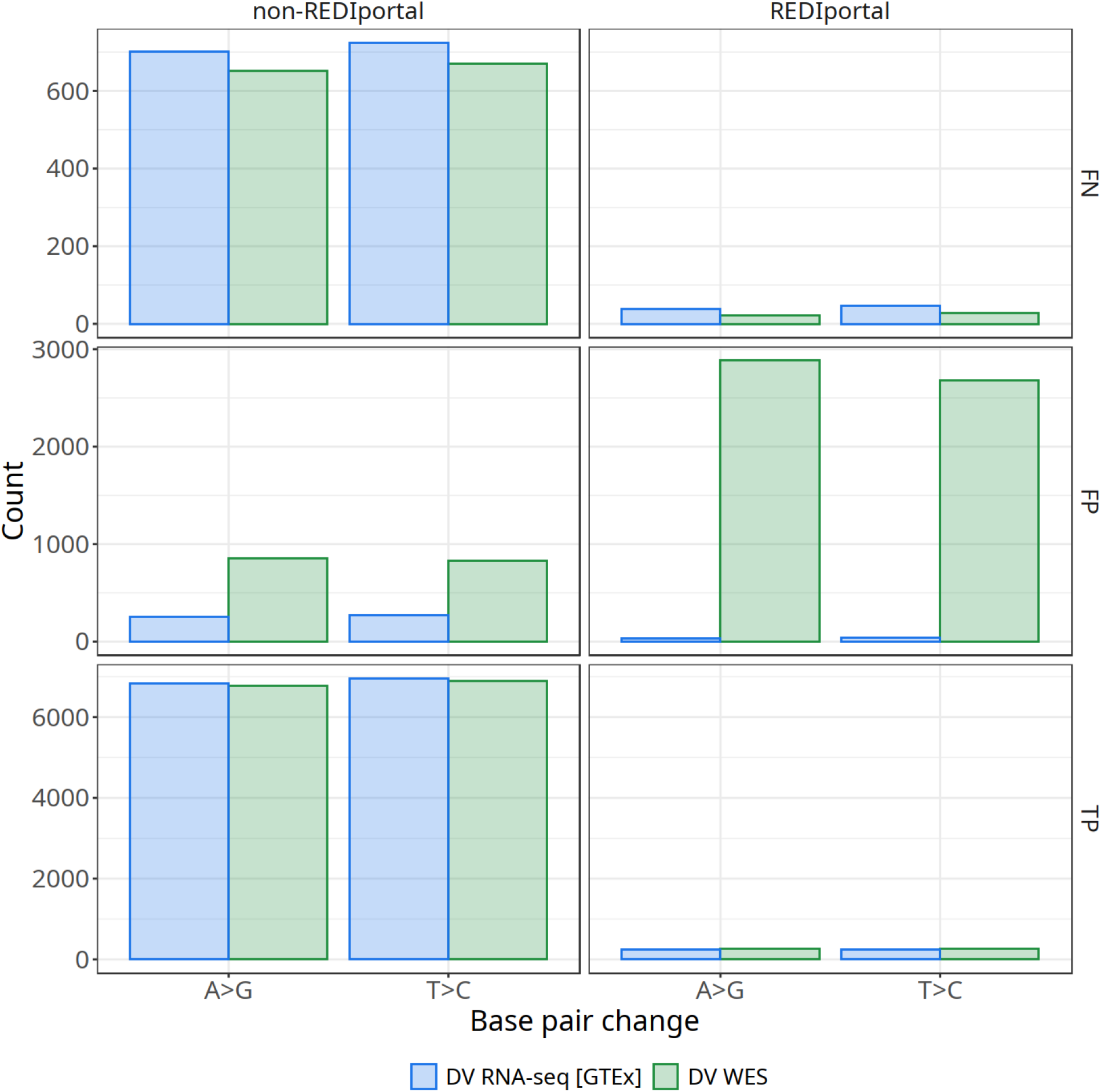
RNA-editing events are ignored. Exonic variants classified as false-negative (FN), false-positive (FP), or true-positive (TP) are shown by base pair change and whether they are present within the REDIportal database for GTEx sample GTEX-ZZ64-1126-SM-5GZXY.

### Selecting a cut-off for practical implementation

When implementing a variant caller into a production analysis pipeline, it is convenient to select a cutoff based on quality scores to maintain a desired true positive (TPR) or false discovery rate (FDR) based on the application. For example, the 1000 Genomes Project FDR target was 5% (although ultimately the project delivered calls at 1% FDR)^36^.

Variability between samples is expected due to both random sampling during the sequencing process and variation among samples. Therefore, cutoff scores should be established by examining multiple replicates or samples using validated variants. We examined 3 HG002 cell-lines (GM24385, GM26105, GM27730), which were evaluated with the DV RNA-seq [GTEx] model in high-confidence GIAB CDS regions to identify an appropriate cut-off score that would maintain a FDR of 1.5% while retaining high sensitivity.

We considered two quality scores to use for setting a cutoff: Genotype quality (GQ) and quality (QUAL). GQ describes the probability of a variant call being incorrect, whereas QUAL describes the probability a given site is not a variant. We found GQ worked well for both SNPs and INDELs for thresholding in comparison to QUAL scores, and decided on a cutoff of GQ>=18 (GQ:**Figure 5**: QUAL: **Supplementary Figure S6**). This threshold achieves a FDR of <1.5% for both SNPs and INDELs while maintaining high sensitivity for SNPs, and modest sensitivity INDELs.

**Figure 5:**
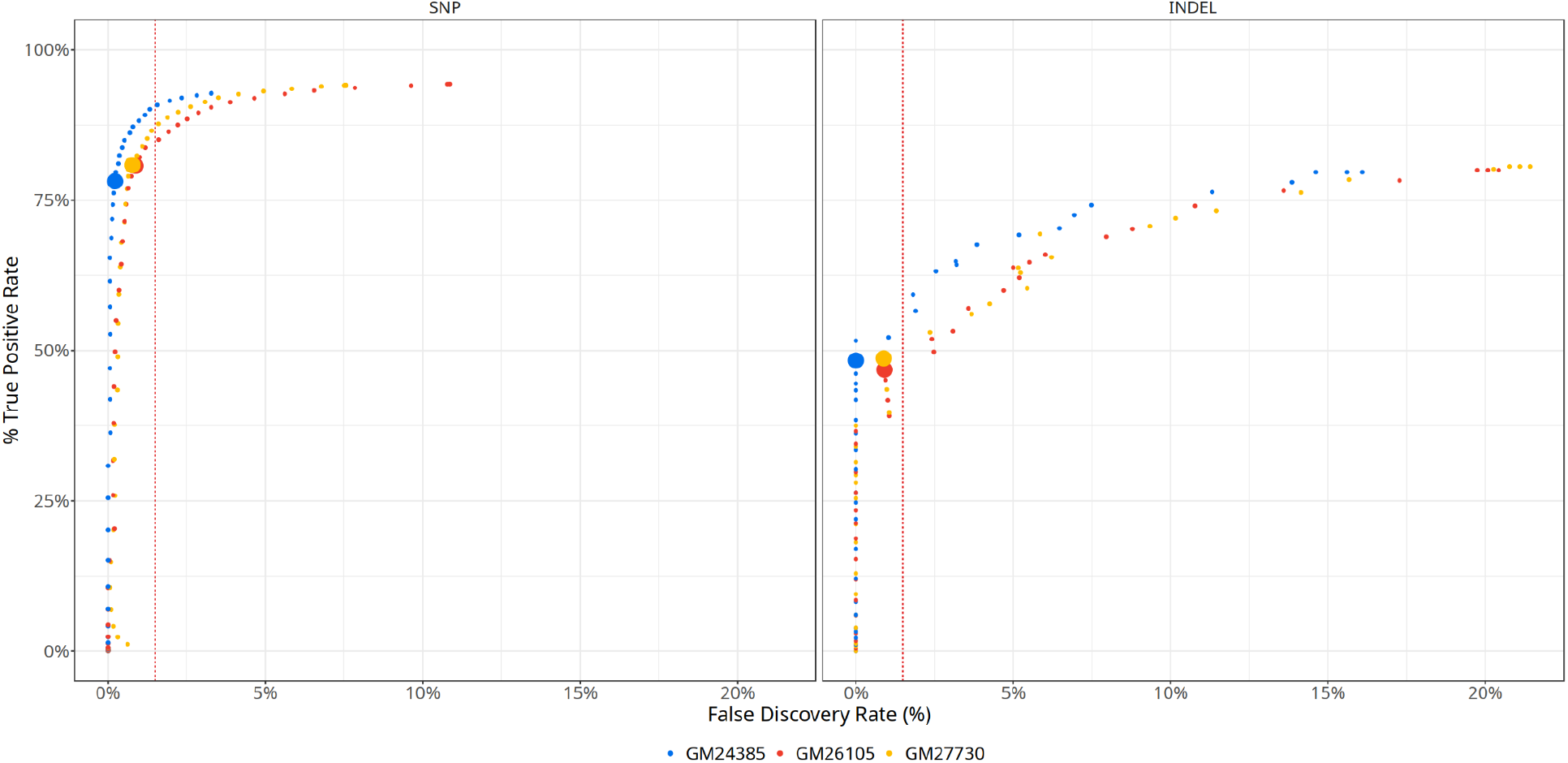
Selection of a QUAL cut-off to maintain FDR of 1.5%. The FDR (x-axis) is plotted against the TPR (y-axis) for both SNP and INDELs across 3 HG002 cell lines in high confidence GIAB regions. Each point represents a given GQ cutoff. The red line indicates a FDR of 1.5%. Larger markers indicate the selected cutoff of GQ>=18.

Finally, we evaluated the GQ>=18 threshold using HG005 (GM26107; poly-A selected) to see how it would perform against unfìltered models and other RNA-seq variant callers. We again examined performance in high-confìdence GIAB regions intersected with CDS regions. We find that DV RNA-seq [GTEx] GQ>=18 is able to achieve a precision of 0.998 on SNPs and 0.989 on INDELs while retaining a high level of sensitivity (**Table 3**).

**Table 3:** Precision and recall in high confidence CDS regions. The DeepVariant RNA-seq [GTEx] model with a GQ>=18 cutoff is listed with alternative RNA-seq variant calling methods. For each variant calling method, we show the precision, recall, F1-score, number of false positives (# FP), and number of true positives (# TP) in high-confidence GIAB regions intersected with CDS regions. Bolded values indicate the best performing variant caller for each column, grouped by variant type (SNP or INDEL).

| Variant Caller | Precision | Recall | F1 | # FP | # TP |
| --- | --- | --- | --- | --- | --- |
| **SNP** |  |  |  |  |  |
| DV RNA-seq [GTEx] GQ>=18 | **0.998** | 0.809 | 0.893 | **18** | 9077 |
| DV RNA-seq [GTEx] | 0.978 | 0.942 | **0.96** | 236 | 10579 |
| Platypus (Opossum) | 0.96 | 0.93 | 0.945 | 431 | 10438 |
| GATK | 0.959 | 0.919 | 0.938 | 444 | 10301 |
| DV WES | 0.938 | **0.946** | 0.942 | 702 | **10624** |
| **INDEL** |  |  |  |  |  |
| DV RNA-seq [GTEx] GQ>=18 | **0.989** | 0.509 | 0.672 | **1** | 86 |
| DV RNA-seq [GTEx] | 0.831 | 0.805 | **0.818** | 28 | 138 |
| Platypus (Opossum) | 0.696 | 0.604 | 0.646 | 45 | 103 |
| GATK | 0.404 | 0.805 | 0.538 | 220 | **149** |
| DV WES | 0.223 | **0.828** | 0.352 | 487 | 140 |

### Runtime

We Examined the runtime performance of our DeepVariant RNA-seq models against existing approaches (**Supplementary Figure 8**). All runtimes were measured using GTEx RNA-seq BAM files on Google Cloud n1-standard-64 virtual machines (64 cores, 240 Gb ram). Using Opossum preprocessed bams generally reduced the required runtime of variant callers, but would add an additional runtime of ~15.6 minutes to preprocess BAM files. DeepVariant RNA-seq models are faster than GATK (**Table 4**). Although Platypus runs faster than DeepVariant, running Platypus requires preprocessing with Opossum. The total runtime of Platypus + Opossum results in a similar runtime to DeepVariant RNA-seq models without Opossum preprocessing (**Table 4**).

**Table 4:** Runtime Performance Of Variant Callers and Opossum on GTEx BAMs.

| model | Duration (Opossum) | Duration (Caller) | Duration (Total) |
| --- | --- | --- | --- |
| Platypus (Opossum) | 932s (~15.53 minutes) | 479s (~7.98 minutes) | 1410s (~23.5 minutes) |
| DV RNA-seq [GTEx] |  | 1426s (~23.77 minutes) | 1426s (~23.77 minutes) |
| DV RNA-seq [GIAB] |  | 1427s (~23.78 minutes) | 1427s (~23.78 minutes) |
| DV WES |  | 1434s (~23.9 minutes) | 1434s (~23.9 minutes) |
| DV WES (Opossum) | 937s (~15.62 minutes) | 1006s (~16.77 minutes) | 1943s (~32.38 minutes) |
| DV RNA-seq [GTEx] (Opossum) | 937s (~15.62 minutes) | 1011s (~16.85 minutes) | 1948s (~32.47 minutes) |
| DV RNA-seq [GIAB] (Opossum) | 937s (~15.62 minutes) | 1017s (~16.95 minutes) | 1954s (~32.57 minutes) |
| GATK |  | 2868s (~47.8 minutes) | 2868s (~47.8 minutes) |

## Discussion

This study introduces DeepVariant RNA-seq, an extension of DeepVariant that enables germline variant calling using Illumina short-read RNA-seq data. To the best of our knowledge, this is the first deep-learning based RNA-seq germline variant caller. However, an existing tool called SmartRNASeqCaller^37^ is a machine-learning based post-processing pipeline that operates on variants called with germline RNA-seq variant callers such as GATK. DeepVariant RNA-seq can operate in a standalone manner, without requiring pre or post processing.

Additionally, we have demonstrated that a “silver-truth set” can be used to generate highly accurate models when used appropriately. The GTEx-based model was trained using label variants derived from the DeepVariant whole-genome sequencing (DV WGS). Although these labels are likely of poorer quality than GIAB-based truth sets, we observed that this approach allowed for a substantially larger and more diverse training dataset than the GIAB-based model. This approach ultimately yielded improved performance compared with the GIAB-based training approach, suggesting that a silver-truth may be appropriate in cases which can dramatically expand gold standard datasets of limited size.

Our examination of RNA editing events suggests our model learns to ignore these types of events, because it is trained on germline variant calls only. Conversely, this suggests that we could train a model capable of calling these RNA-editing events by configuring our model to produce an additional output (*e.g*. probability of a site being an RNA editing event). Such functionality could integrate germline and RNA-editing events into a single variant call format (VCF) file, and allow for investigations of RNA editing and germline variant interactions. Consider the result when a heterozygous variant occurs adjacent to an RNA editing event within a codon. For example, [heterozygous genotype: T/C] + [RNA edit: A/G] + [C] could produce four different protein isoforms: UAC (Tyr), UGC (Cys), CAC (His), and CGC (Arg). Although CDS editing is rare, this extension of our work would allow for these types of events to be identified, in addition to long-range combinatorial effects of genotype and RNA edits.

By releasing the DeepVariant RNA-seq models, we hope to provide reliable methods to add RNA-seq variant detection to typical RNA analysis use cases. We welcome community feedback to further enhance DeepVariant RNA-seq.

## Availability of Software and Data

The DeepVariant RNA-seq GTEx model and documentation are available at https://github.com/google/deepvariant.

Supplementary Tables are available [here](https://docs.google.com/spreadsheets/d/1B8PJvTiSu3VulUmX0a-MxZ7E2VNme8-PCxmPrbGkTas/edit?usp=sharing&resourcekey=0-q2frxtotnvpbXDfbdQO-Wg).

URLs for GIAB RNA-seq FASTQs and BAM are available and are listed in **Supplementary Table 4**.

## Supplementary Figures and Tables

### Supplementary Tables

**Supplementary Table S1**-GTEx sample information.

**Supplementary Table S2**-Summary of GTEx and GIAB datasets.

**Supplementary Table S3**-Performance across variant callers, restricted to chr20 only.

**Supplementary Table S4**-URLs for GIAB RNA-seq FASTQs and BAM Files.

### Supplementary Figures

**Supplementary Figure S1:**
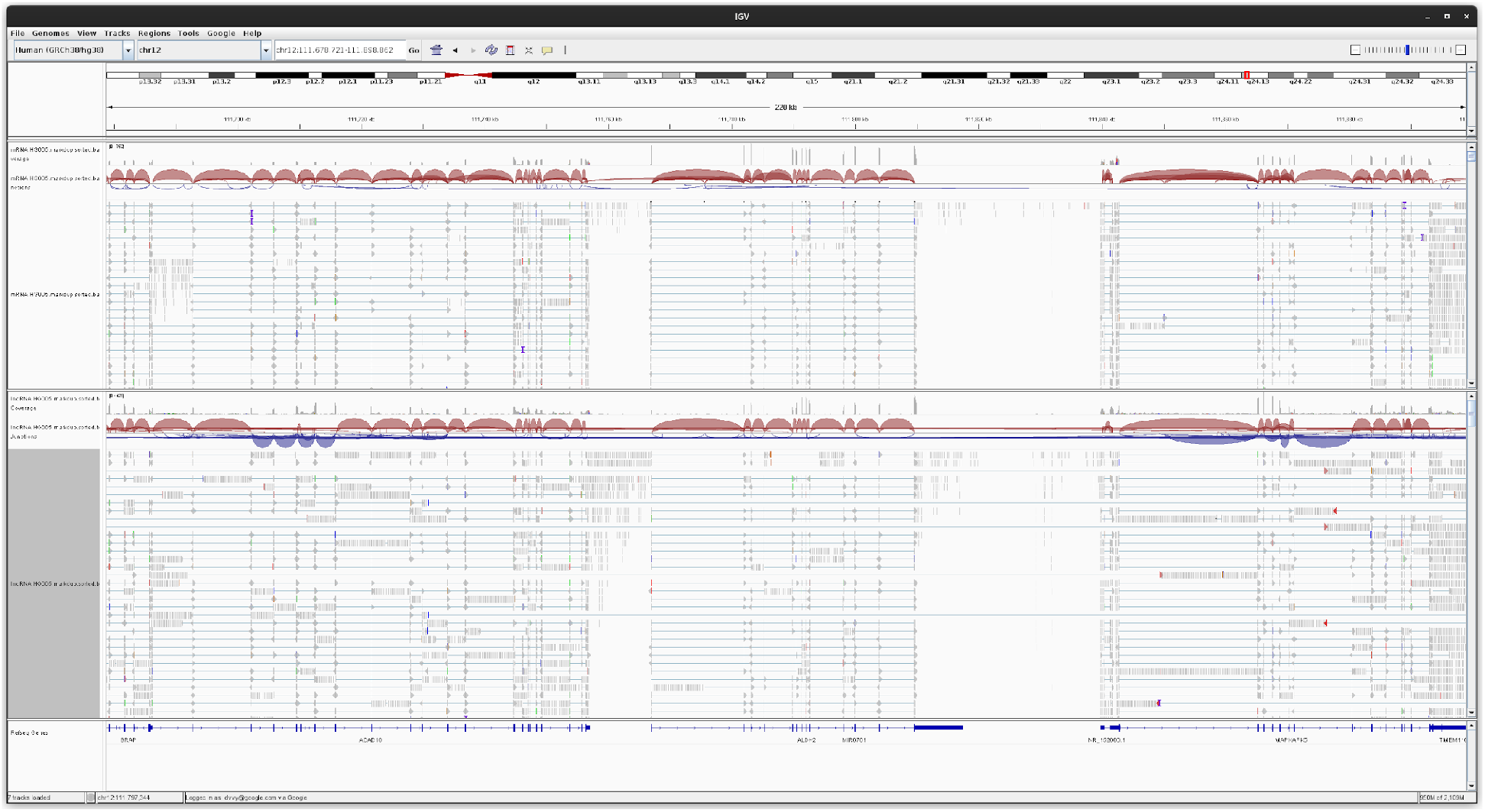
RNA-seq data. A screenshot of the Integrative Genomics Viewer (IGV) showing skip regions present in RNA-seq data.

**Supplementary Figure S2:**
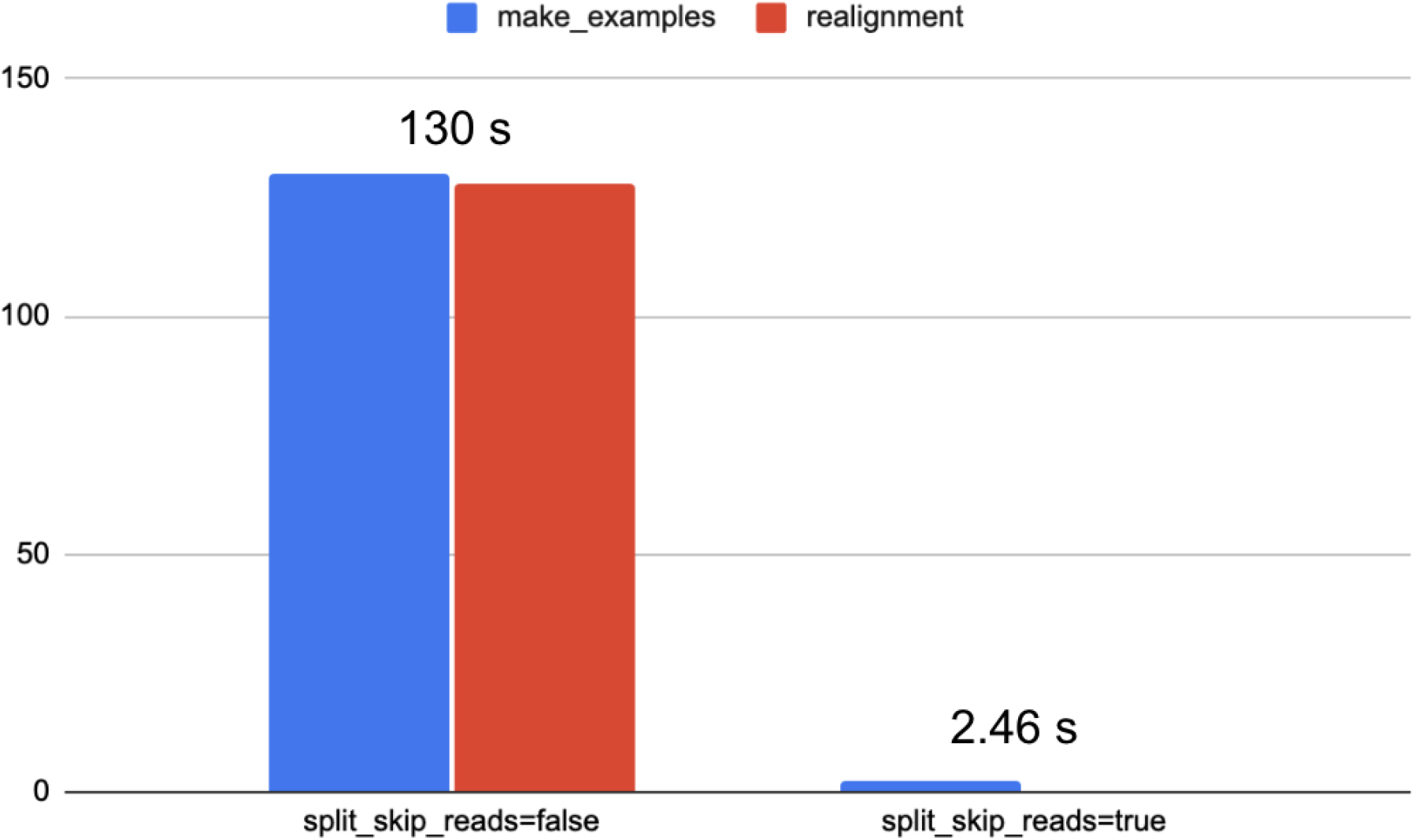
The split_skip_reads flag. The time required to make examples and perform realignment is shown in seconds for chromosome 1:1-100kb. Setting `--split_skip_reads` significantly reduces the time required to perform local-realignment when processing RNA-seq data.

**Supplementary Figure S3:**
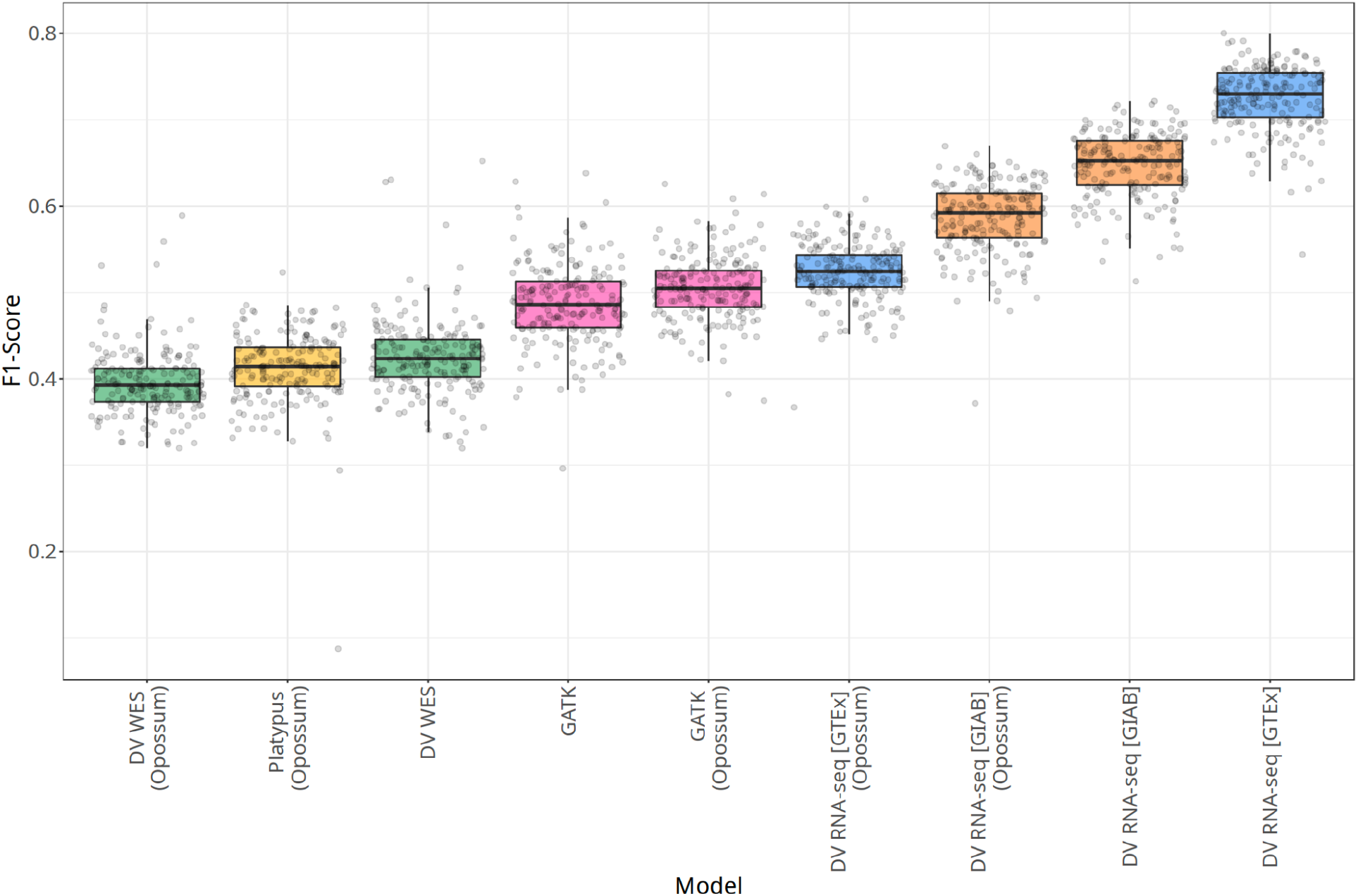
INDEL F1-scores in CDS regions for 200 RNA-seq samples. INDEL F1 scores in CDS regions are shown for 200 GTEx RNA-seq samples across Platypus, GATK, DeepVariant WES, and DeepVariant RNA-seq. Data that was pre-processed with Opossum is labeled as such.

**Supplementary Figure S4:**
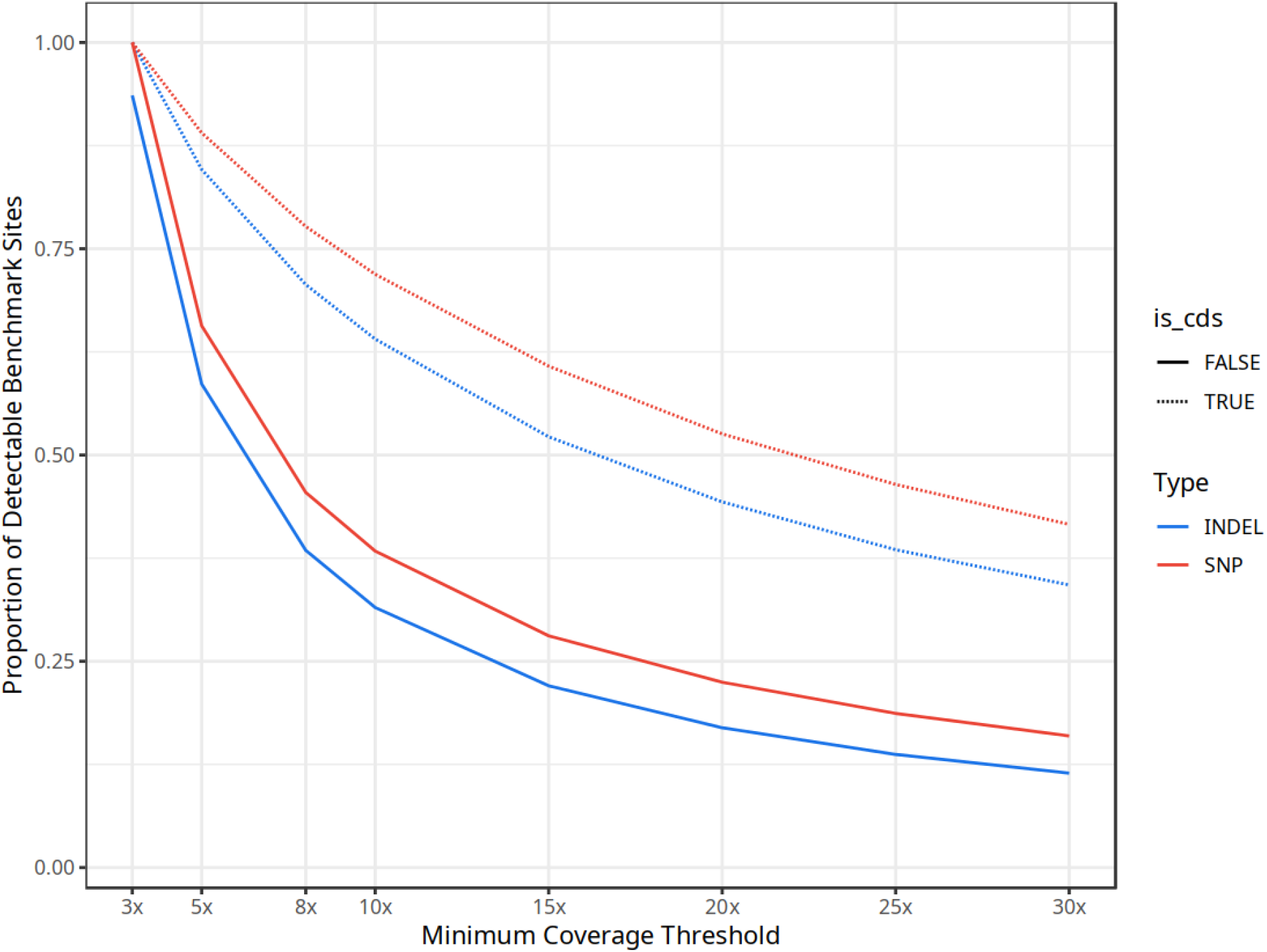
Proportion of detectable sites. The proportion of detectable benchmark sites are shown across minimum coverage thresholds. “Detectable” sites are label sites present within 3x regions that remain at higher coverage thresholds. Curves are shown for SNP and INDEL variants present within or outside of CDS regions. These curves show how the proportion of 3x variant sites is reduced when imposing higher minimum coverage thresholds.

**Supplementary Figure S5:**
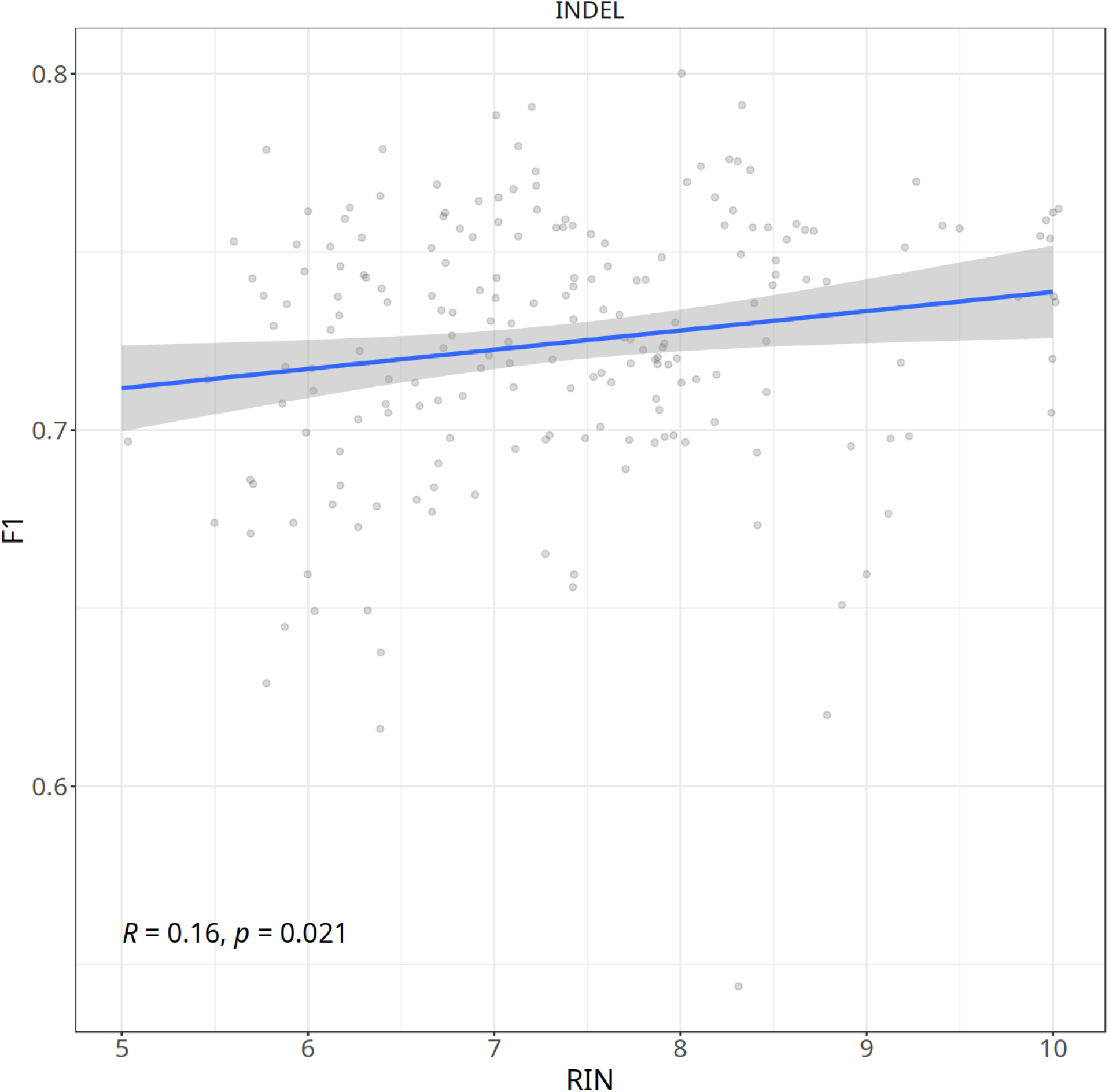
Correlation between RIN and F1-score for INDELs in CDS regions. The correlation between RIN and the F1 score is shown for 200 RNA-seq samples for INDEL calls in CDS regions.

**Supplementary Figure S6:**
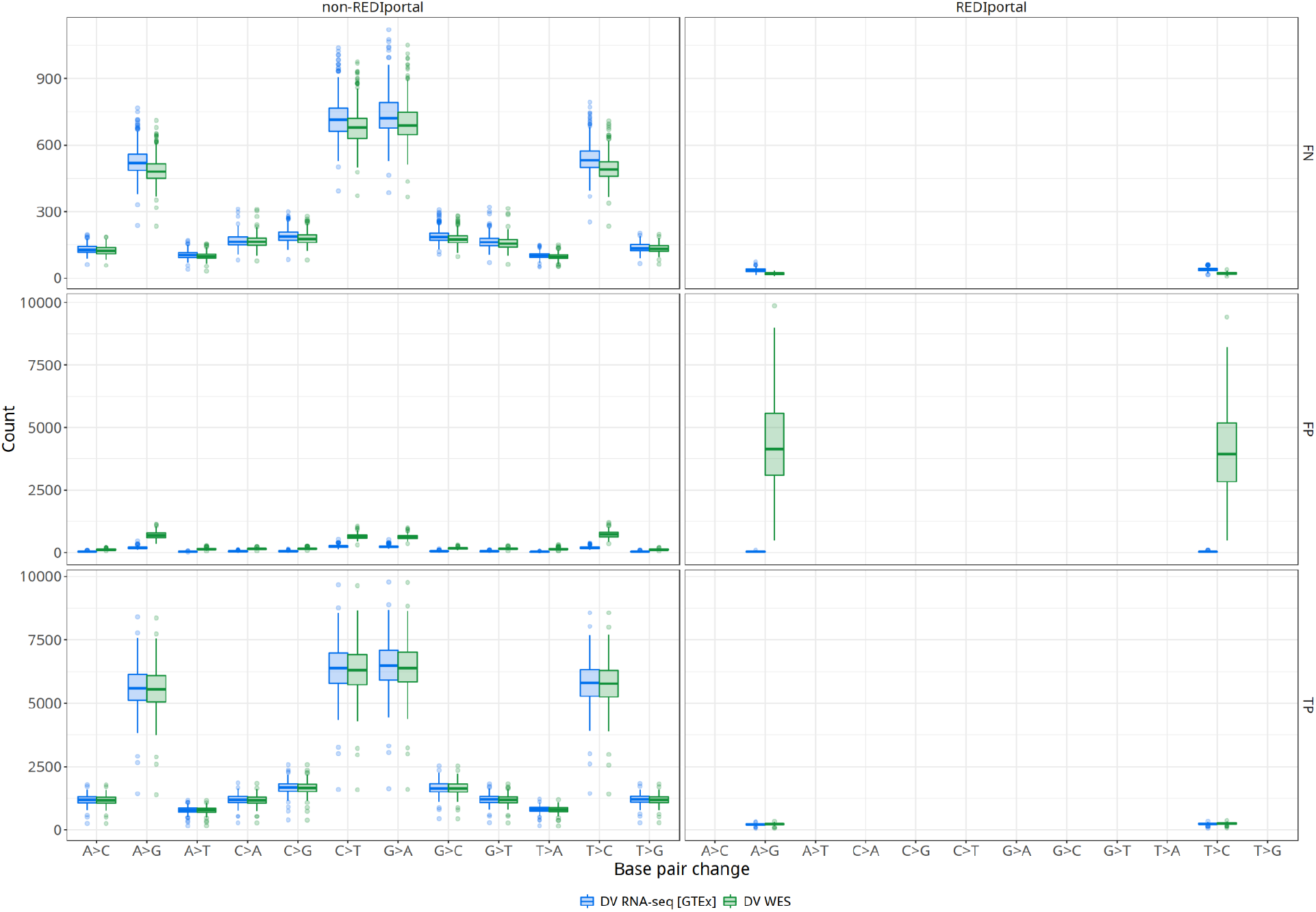
Aggregate summary of RNA edited site classification. The distribution of false negative (FN), false positive (FP) and true positive (TP) events is shown by base change and whether or not a variant has previously been characterized as an RNA edit event in the REDIportal database. Results are shown for the DeepVariant RNA-seq and DeepVariant WES models.

**Supplementary Figure S7:**
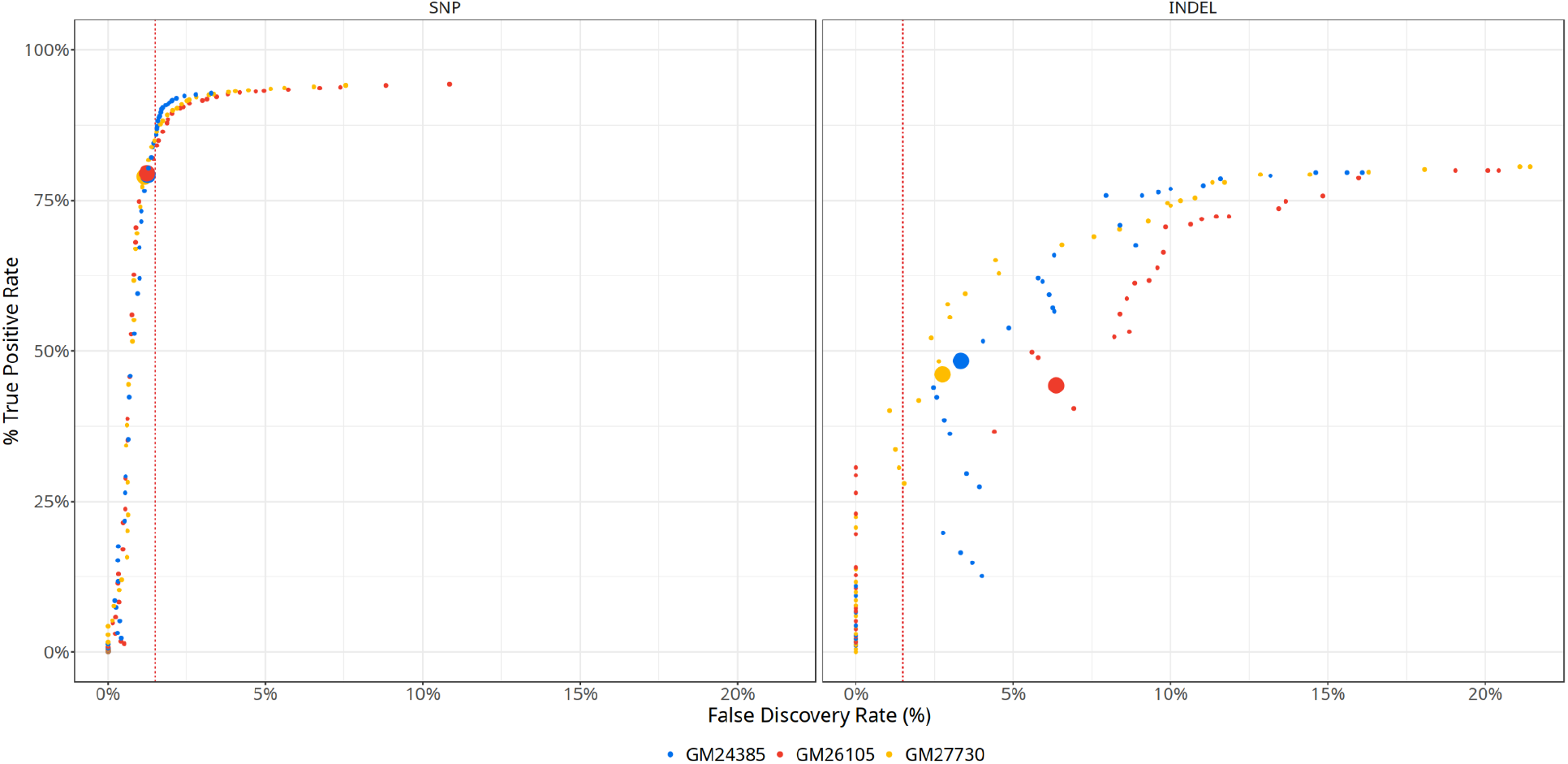
Selection of a QUAL cut-off to maintain FDR of 1.5%. The FDR (x-axis) is plotted against the TPR (y-axis) for both SNP and INDELs across 3 HG002 cell lines. Each point corresponds with the FDR (x-axis) and TPR (y-axis) at a given QUAL cutoff. The red line indicates a FDR of 1.5%. Larger markers indicate a cutoff of QUAL>=25.

**Supplementary Figure S8:**
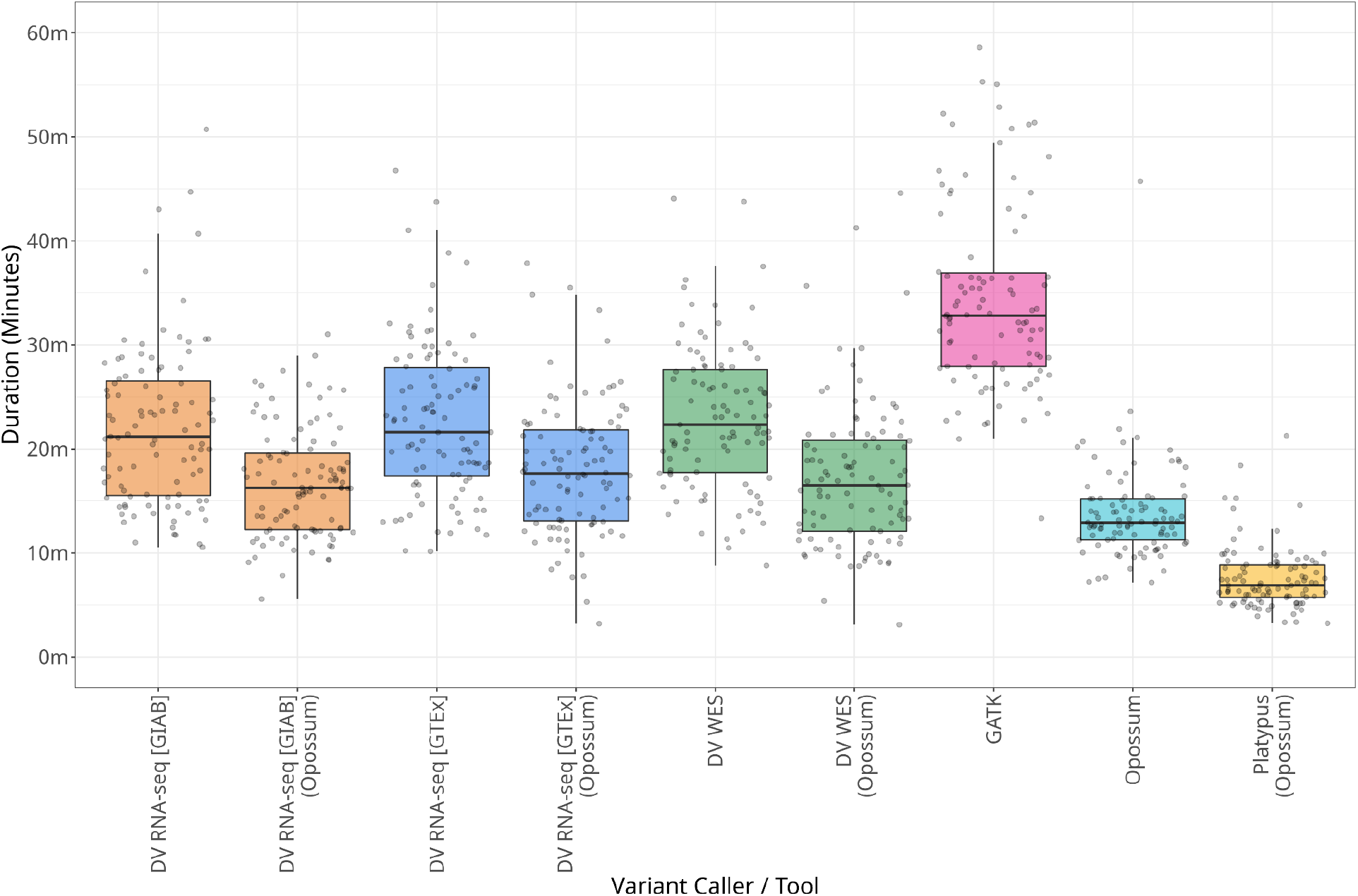
The runtime distribution of a random sample of 100 runs for each variant caller and Opossum are shown. Filtering was performed to remove excessive runtime outliers (>1 hr). These were only observed for GATK.

